# Easyml: Easily Build and Evaluate Machine Learning Models

**DOI:** 10.1101/137240

**Authors:** Woo-Young Ahn, Paul Hendricks, Nathaniel Haines

**Affiliations:** Department of Psychology Seoul National University Seoul, Korea 08826; Department of Psychology The Ohio State University Columbus, OH 43210, USA

**Keywords:** machine learning, data science, supervised learning, data mining, visualization, R, Python

## Abstract

The `easyml` (easy <u>m</u>achine <u>l</u>earning) package lowers the barrier to entry to machine learning and is ideal for undergraduate/graduate students, and practitioners who want to quickly apply machine learning algorithms to their research without having to worry about the best practices of implementing each algorithm. The package provides standardized recipes for regression and classification algorithms in R and Python and implements them in a functional, modular, and extensible framework. This package currently implements recipes for several common machine learning algorithms (e.g., penalized linear models, random forests, and support vector machines) and provides a unified interface to each one. Importantly, users can run and evaluate each machine learning algorithm *with a single line of coding.* Each recipe is robust, implements best practices specific to each algorithm, and generates a report with details about the model, its performance, as well as journal-quality visualizations. The package’s functional, modular, and extensible framework also allows researchers and more advanced users to easily implement new recipes for other algorithms.

## 1. Introduction

Numerous machine learning libraries (or packages) are becoming available in popular programming languages, especially R (R Core Team, 2016) and Python (Rossum, 1995). Both languages are high-level, interpreted, employ functional and object-oriented paradigms, and have a wide ecosystem of mature machine learning libraries. However, existing machine learning libraries assume the user has a solid understanding of statistics and machine learning principles and best practices, strong programming skills, and the knowledge of how to apply this skillset to their problem. Oftentimes, this is not the case. Individuals without strong technical background increasingly want to apply machine learning techniques to their research without having to spend years studying mathematics, statistics, and/or computer science and there is a critical need to lower the barrier to machine learning or computational approaches in general (Ahn and Busemeyer, 2016; Ahn et al., 2017). The `easyml` targets these individuals and hopes to lower the barrier to entry to machine learning by providing user-friendly recipes for common machine learning algorithms.

These recipes leverage R and Python’s programming capabilities and their existing machine learning libraries and breaks down each analysis into steps common to all algorithms and steps unique to each algorithm. These steps are abstracted from the user by a common unified framework. Thus, machine learning is like baking a cake; whether one wants to bake a chocolate cake or a vanilla cake, one still needs eggs, flour, and butter as the core ingredients. If you mix them in certain steps and add chocolate, a chocolate cake is baked. If you mix the core ingredients in a certain way and add vanilla, a vanilla cake is baked. Analogously, though one may run similar steps to build and evaluate a penalized linear model (Friedman et al., 2010; Simon et al., 2011) and a random forest model (Breiman, 2001), one will wish to assess the coefficients of the linear model and the variable importances of the random forest model. easyml (easy machine learning) makes this easy by handling the best practices for each algorithm but still allows an advanced user the flexibility to customize each recipe.

## 2. Project Vision

### Maintenance

This package is maintained by Paul Hendricks and Woo-Young Ahn.

### Availability

The easyml source code is available under the MIT license and hosted on GitHub (https://github.com/CCS-Lab/easyml).

### Standardized recipes

The package provides standardized recipes for regression and classification machine learning algorithms in R and Python (see Table 1). Specifically, `easyml` provides recipes and unified interface to some of widely used machine learning algorithms including penalized regression models, random forests, support vector machines (Cortes and Vapnik, 1995), Group-Lasso interaction model (Lim and Hastie, 2015), and (deep) neural network models. More advanced users will find it easy to implement new recipes for other algorithms. To implement the algorithms, we use other R and Python packages including `glmnet` (Friedman et al., 2010), `randomForest` (Liaw and Wiener, 2002), `e1071` (Meyer et al., 2017), `glinternet` (Lim and Hastie, 2015), `nnet` (Venables and Ripley, 2002), `darch` (Drees, 2013), and `scikit-learn` (Pedregosa et al., 2011). We also plan to add more algorithms in the future.

**Table 1.**
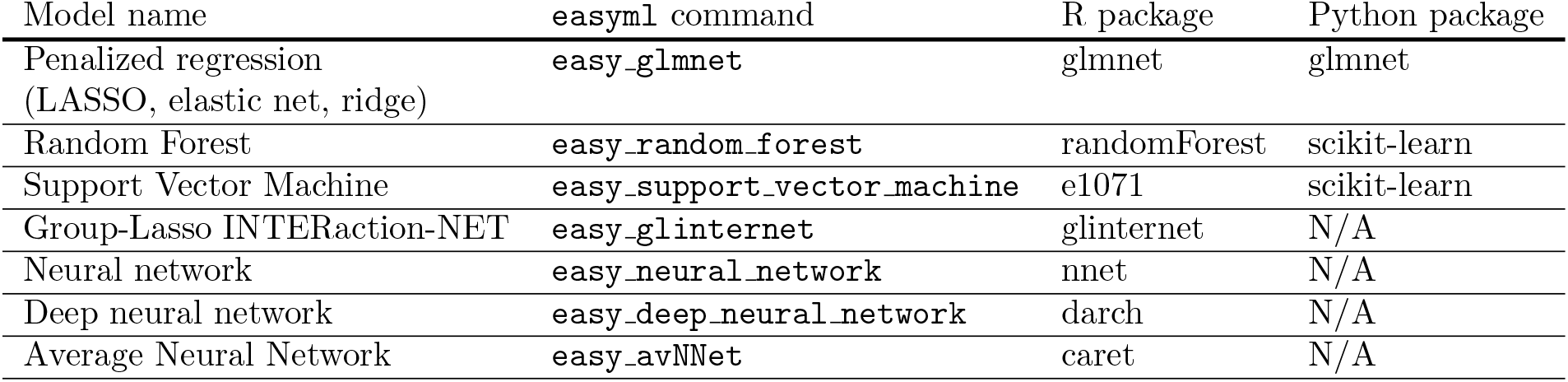
List of machine learning models currently implemented in easyml (as of v0.1.0).

### Journal-quality visualizations

Users will find that `easyml` can immediately produces journal-quality visualizations. These visualizations can be easily be modified, if needed, and used directly in research papers or presentations. See Section 4 for an example.

### Functional, modular, and extensible framework

The package’s functional, modular, and extensible framework also allows researchers and more advanced users to implement new recipes for other algorithms. An example of how to implement a new algorithm is demonstrated in Section 5.

### Parallelization

The nature of machine learning often lends itself to highly parallelizable code. `easyml` makes it possible to run all recipes leveraging as many CPUs as are available. Simply specify the `n_core` parameter in the interface and `easyml` will parallelize the analyze over that number of cores.

### Code quality control

`easyml` uses software engineering best practices such as Continuous Integration (CI) to check the build of the package, Unit Testing and Code Coverage to check the quality of the code, and linting to ensure adherence to a common style. As of this writing, all builds and tests pass on Ubuntu 14.04 and Mac OS X and the Code Coverage is above 85%. The project is also hosted on GitHub (https://github.com/CCS-Lab/easyml), and is available to users who want to examine the source code, contribute to the code base, or provide the authors with feedback or alert the authors to potential bugs, both via issues.

### Documentation

`easyml` provides exhaustive documentation and examples for both R and Python. Users interested in the R package can find documentation here: http://ccs-lab.github.io/easyml. Users interested in the Python package can find documentation here:http://easyml.readthedocs.io.

## 3. Recipes

`easyml` uses standardized recipes for regression and classification machine learning algorithms in R and Python. These recipes can be broken down into multiple steps and are useful for interpreting models (e.g., estimating coefficients and variable importances) or estimating in-sample and out-of-sample performance (e.g., predictions and measures of goodness-of-fit). For each of these recipes, we describe our motivation for including the recipe, a breakdown of the steps in each recipe, and the algorithms that recipe is implemented for.

### Coefficients

Linear models are powerful due to their simplicity, robustness, and interpretability of variables. However, sometimes the estimated coefficients for linear models are different after each run, even with the same random state. This can be due to the low-level code not setting the random state at the C/Fortran level or due to the stochastic nature of the algorithm or optimizer. This phenomenon makes it difficult to interpret a coefficient after building the model only once. To account for this intrinsic randomness and ensure the final coefficients returned are robust estimators, we generate the coefficients `n_samples` times using k-fold cross validation, where `n_samples = 1000` and k = 10 are set as the defaults, and then calculate the mean and standard deviation of the estimated coefficients. We have applied and validated this protocol in previous studies (Ahn et al., 2014; Ahn and Vassileva, 2016; Ahn et al., 2016; Vilares et al., 2017). The ability to generate beta coefficients is currently implemented only for the penalized linear model algorithm (`easy_glmnet`). See Algorithm 1.

#### Algorithm 1 Generate coefficients

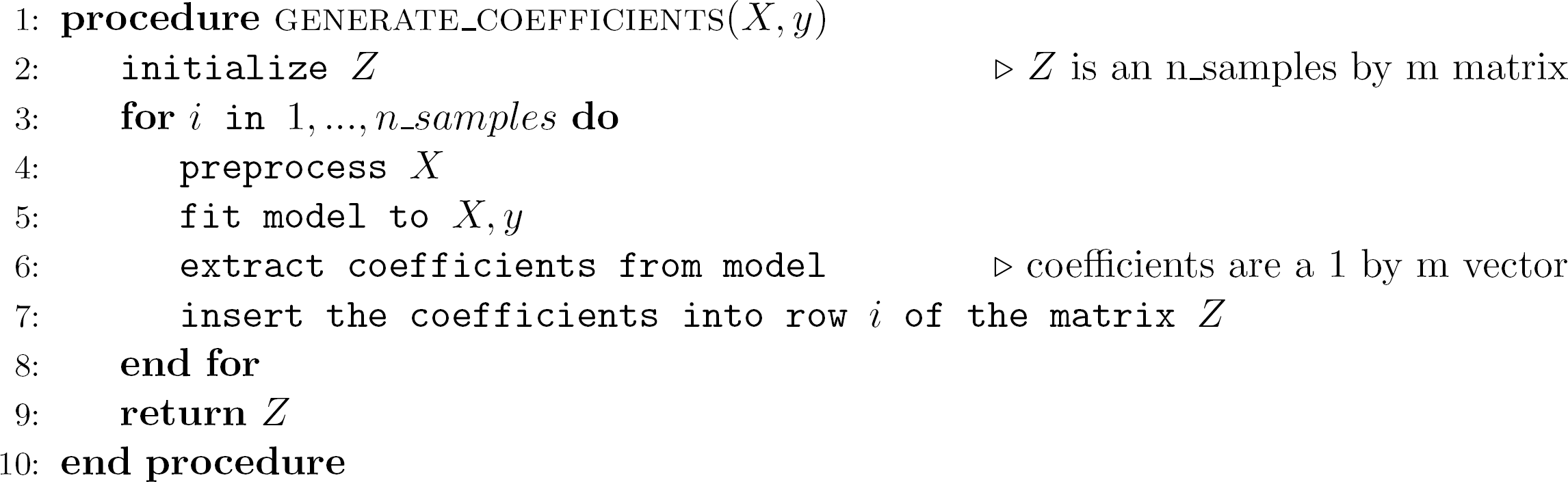

### Variable Importances

Ensemble models are powerful due to their simplicity, ability to capture the non-linear patterns of features from the data, and like linear models, their interpretability of variables. As with linear models, we wish to calculate and visualize the importances of the variables as part of our machine learning protocol. Like linear models, ensemble models often have inherent sources of randomness. For example, the random forest algorithm bootstraps the data randomly and randomly selects a subset of predictors to use in each decision tree. Interpretability heuristics such as variable importance scores can often differ from one random state to another. To ensure the resulting variable importances are robust, we can generate the random forest algorithm `n_samples` times, where `n_samples = 1000` is set as the default, and then calculate the mean and standard deviation of the estimated importances. The ability to generate variable importances is currently implemented for the random forest algorithm (`easy_random_forest`), which was used in our recent paper (Haines et al., in preparation). See Algorithm 2.

### Predictions

We often wish to visualize our predictions, whether it’s a plot of actual against predicted values or a plot of the area under the curve (AUC) of a Receiver Operating Characteristic (ROC) curve. If models sometimes produce random, albeit small, deviations in coefficients or weights, these deviations can propagate to our predictions. To guard against this intrinsic error, we train a model using k-fold cross validation within the training set (k = 10 set as the default) and generate predictions `n_samples` times for a particular train-test split (separately on training and test sets). Then we average the predictions across the `n_samples` iterations. By default, the training and test sets are 67% and 33% of the whole dataset and it can be adjusted (e.g., `train_size = 0.67`). The ability to generate predictions is currently implemented for all algorithms. See Algorithm 3 (note that *nrow*(*X*_*z*_) indicates the number of rows in *X*_*z*_).

#### Algorithm 2 Generate variable importances

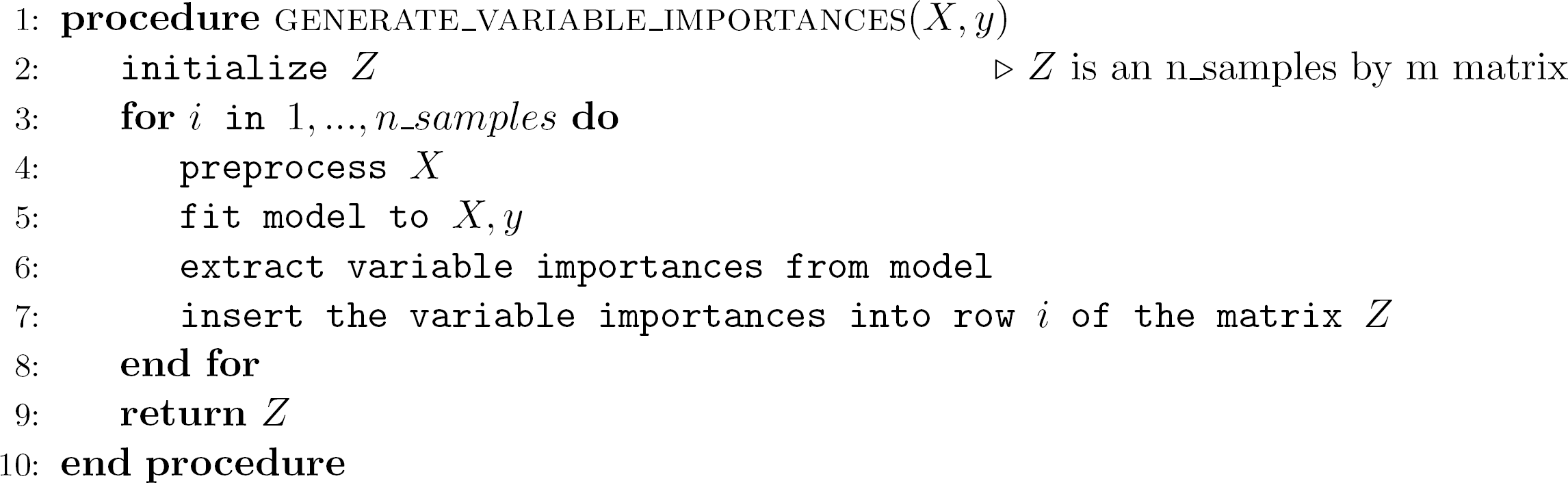

#### Algorithm 3 Generate predictions for single train-test split

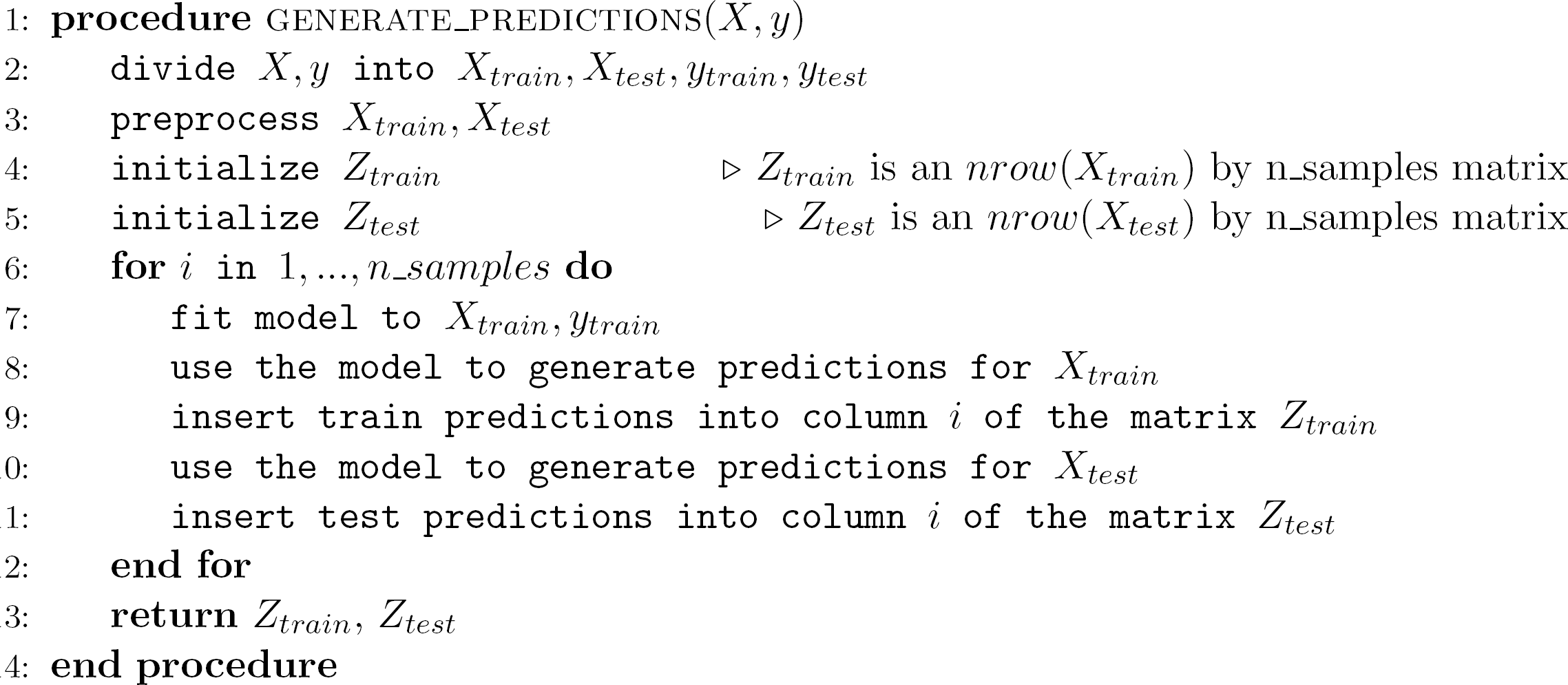

### Model performance

Often we wish to visualize *model performance* representing the quality (i.e., accuracy) of our predictions, whether it’s a plot of mean squared errors, correlation coefficients, or AUCs. We can guard against intrinsic errors by replicating predictions many times for a particular train-test split, averaging the predictions across `n_iterations`, generating a model performance metric, and replicating for many (`n_divisions`) different train-test splits. The reader is referred to the Algorithm 4 box for more details. The ability to generate model performance is currently implemented for all algorithms.

## 4. Example

This example demonstrates how to use `easyml` in both R and Python. For further examples on how to use `easyml` in R, please see the documentation at http://ccs-lab.github.io/easyml. For further examples on how to use `easyml` in Python, please see the documentation at http://easyml.readthedocs.io. In this example, we will use `easyml` to replicate findings reported in Ahn et al. (2016) where a penalized logistic regression was used to identify multivariate patterns of behavioral measures that can classify individuals with cocaine dependence. To use `easyml` in R, we must first install the `easyml` library.

### Algorithm 4 Generate model performance metrics

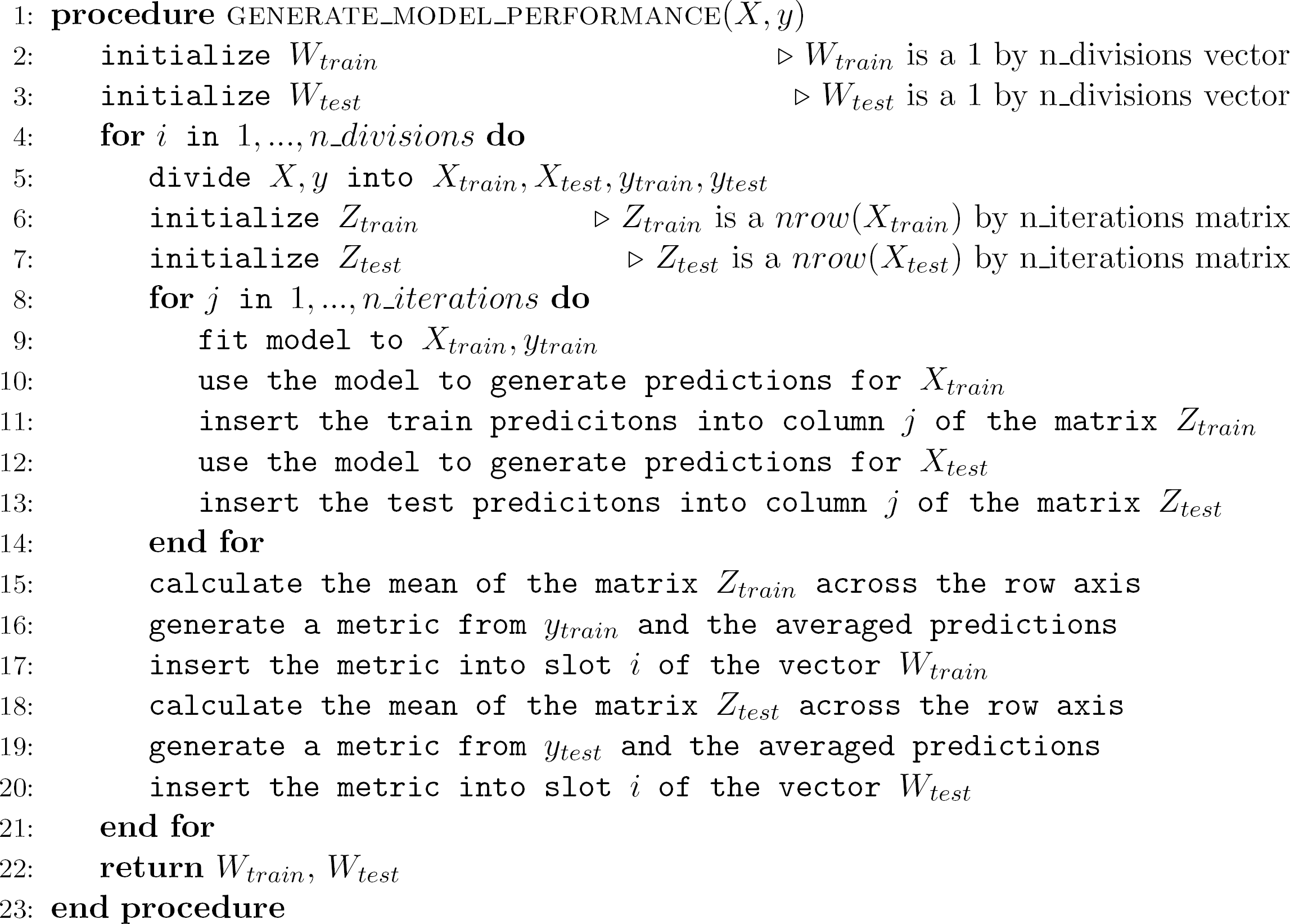

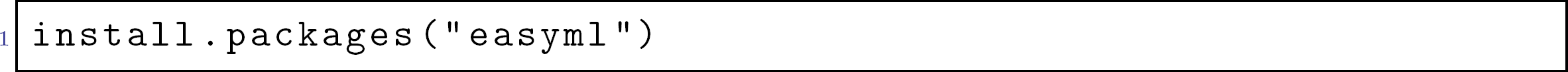

To use `easymlpy` in Python, we must first install the `easymlpy` library using pip.

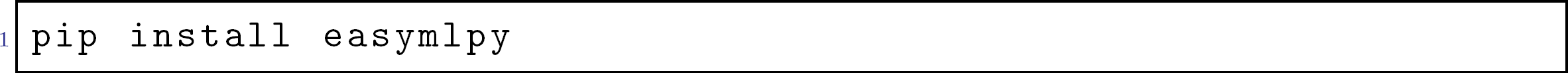

Next, let’s load the package and the data set in R.

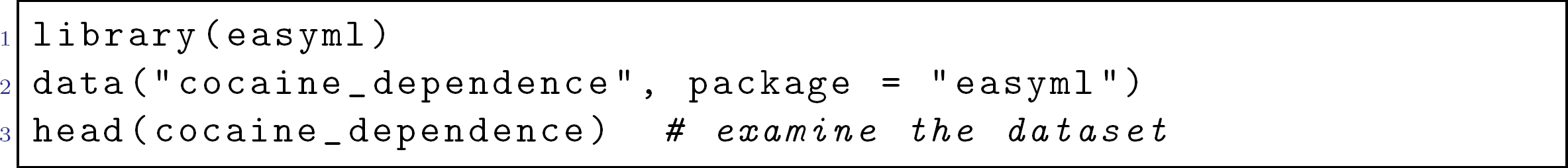

And in Python:

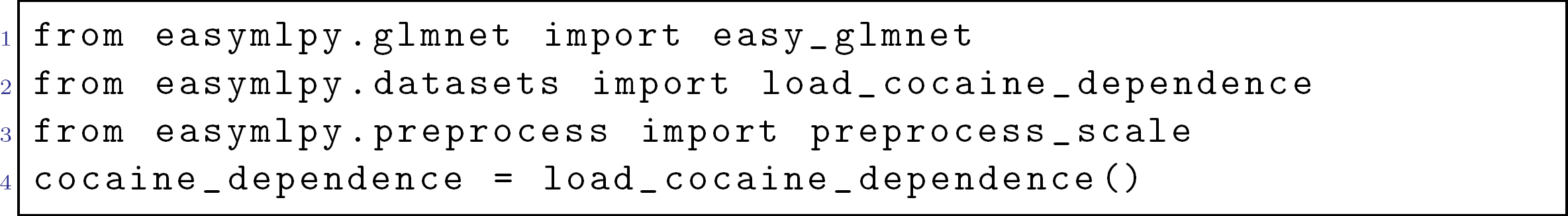

Finally, we pass in the following input arguments to run the analysis:

- `.data`, (or just `data` within Python) the data set to be analyzed (*n* × *m* matrix). *n*=the number of samples. *m*=the number of features. At this time, it should contain no missing data.
- `dependent variable`, the name of the dependent variable, which is an n by 1 vector. In the cocaine data, `diagnosis` (0=healthy control, 1=cocaine user) is the dependent variable.
- `family`, the name of the family of regression with choices are “gaussian” and “binomial”. Since we are modeling a binary dependent variable, we will select “binomial”.
- `preprocess`, the preprocessing function to use on the data. We choose the `preprocess_scale` function so as to scale (z-score) any continuous variable across samples (full data set or train/test data sets) before training a model.
- `exclude_variables`, which variables, if any, should be excluded from the analysis. If there is more than one variable, use the function c() (e.g., `exclude_variables = c` ("subject", "edu_yrs")).
- `categorical_variables`, which variables are categorical, and thus need to be specially handled during preprocessing. Note that categorical variables will not be normalized. If there is more than one variable, use the function `c()`.
- `random state`, the seed to use for the random state.
- `model_args`, the list of arguments specific to penalized linear models. See `?glmnet:: glmnet`.

In R:

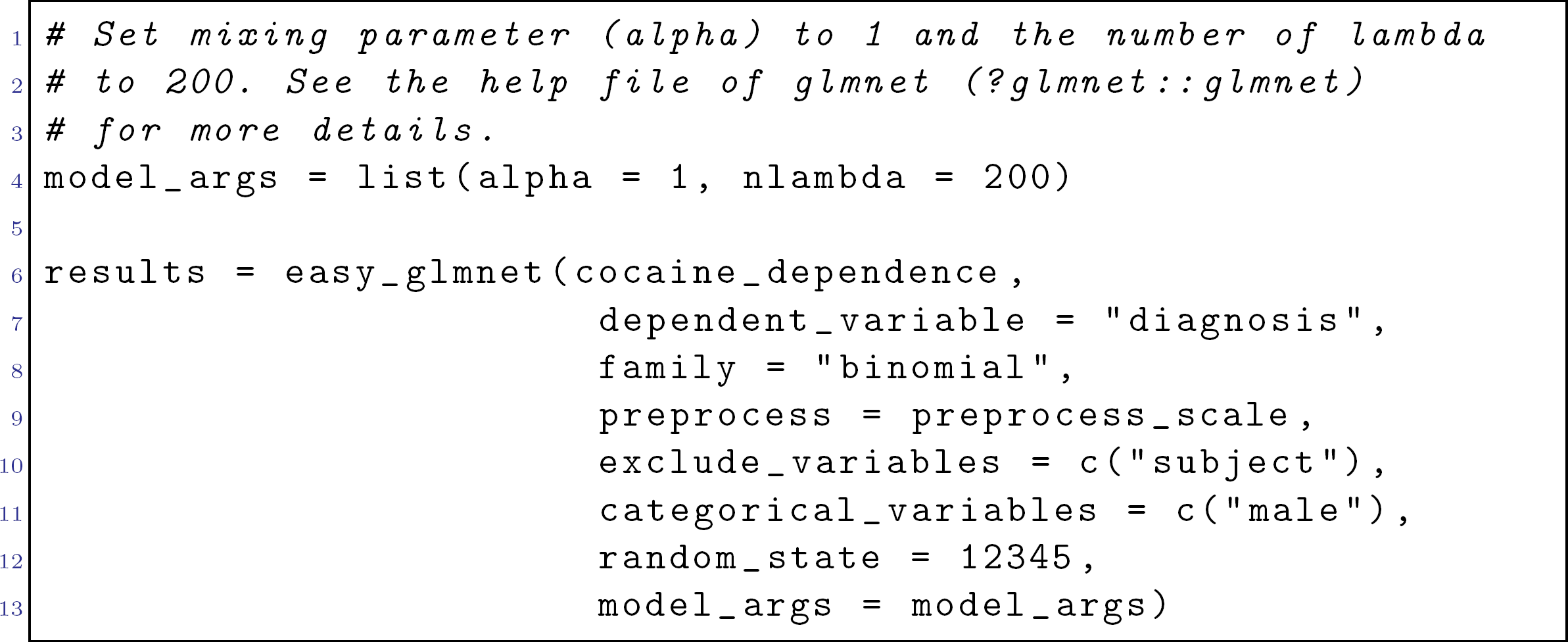

And in Python:

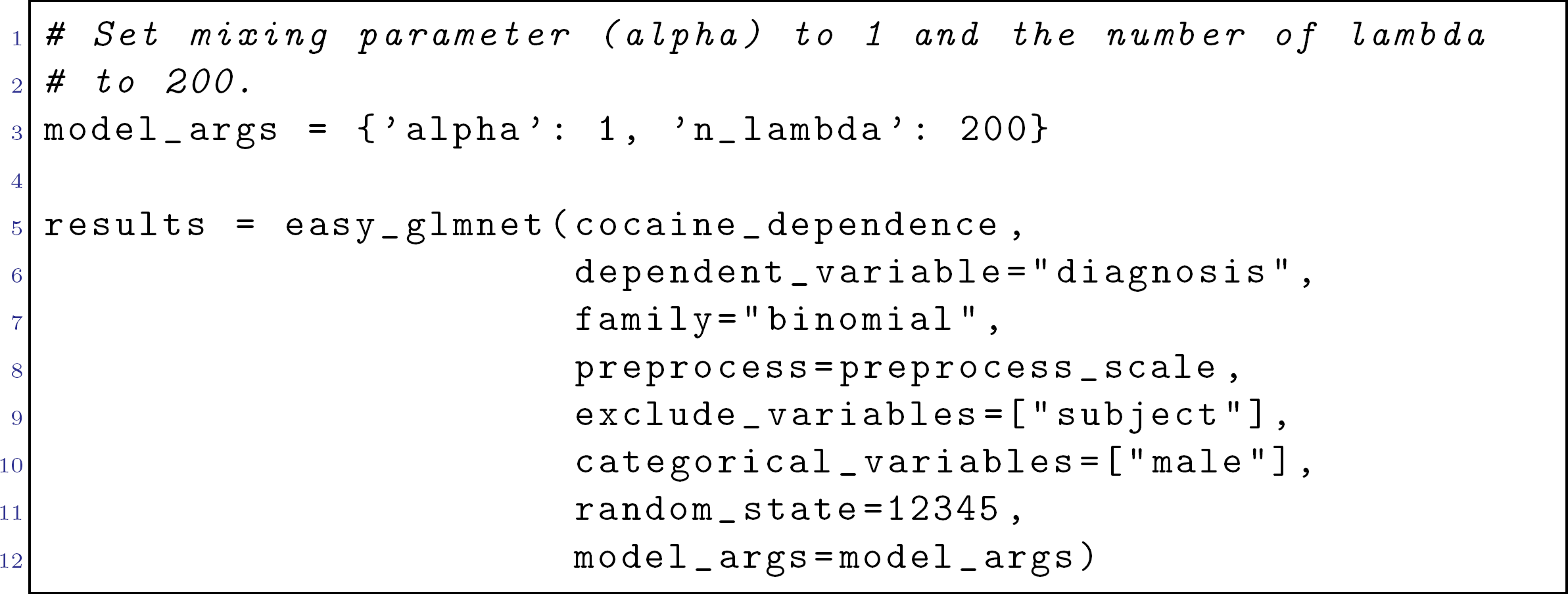

That’s it! Now let’s examine the results. Each algorithm returns a list with objects for various functions, data structures, and plot objects that are instrumental to the analysis. Calling the `names` function in R on the variable `results` will show all the slots available to us.

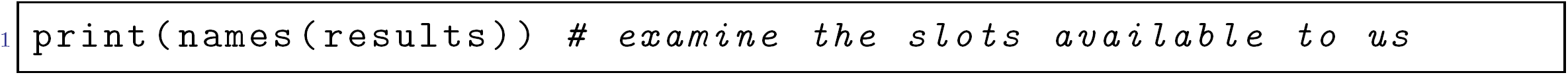

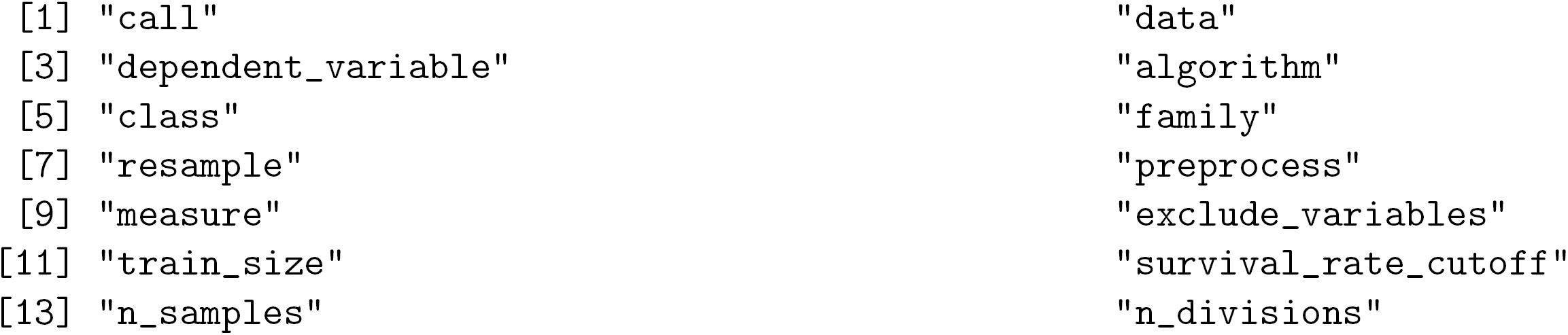
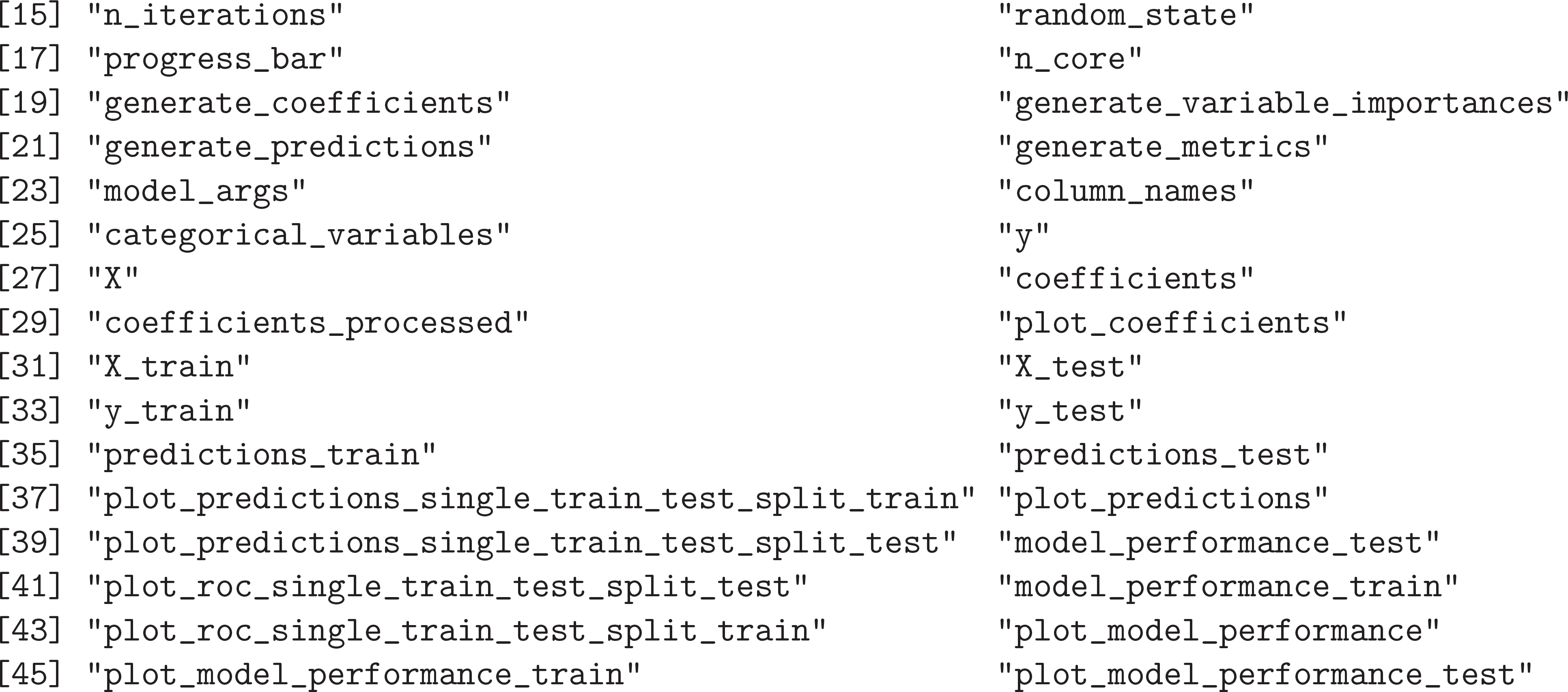

In R, `ggplot2` objects can be accessed via the $ operator. For example, to examine the ROC Curve for the train data set, we can call the following (see Figure 1A):

**Figure 1:**
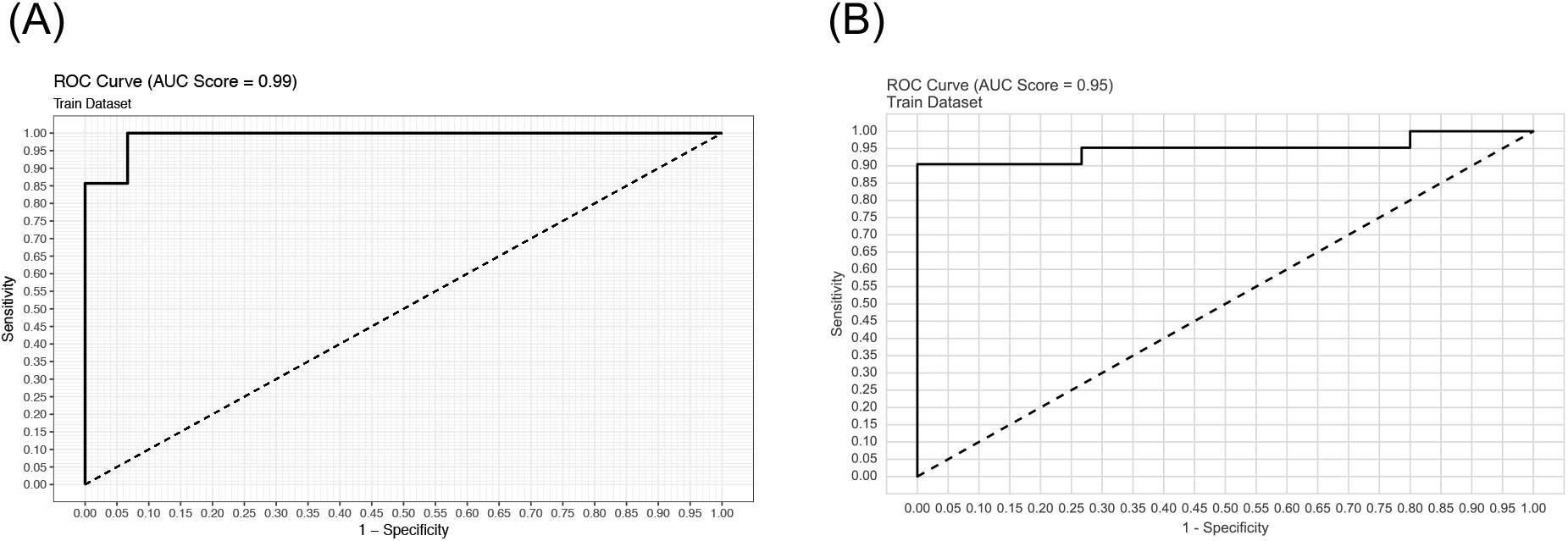
ROC Curve for the train dataset in (A) R and (B) Python.

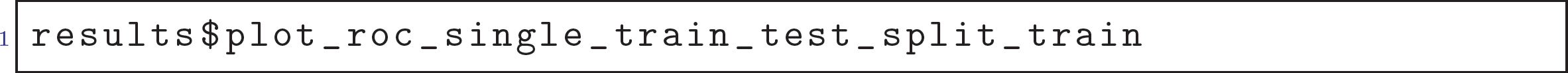

In Python, functions that produce `matplotlib` objects can be accessed via the . operator. To mirror the call in Python, we can call the following (see Figure 1B):

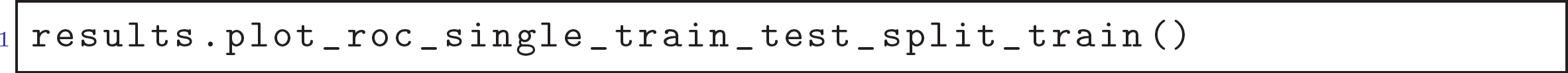

And we can examine the out-of-sample predictions for the test data set (see Figure 2A).

**Figure 2:**
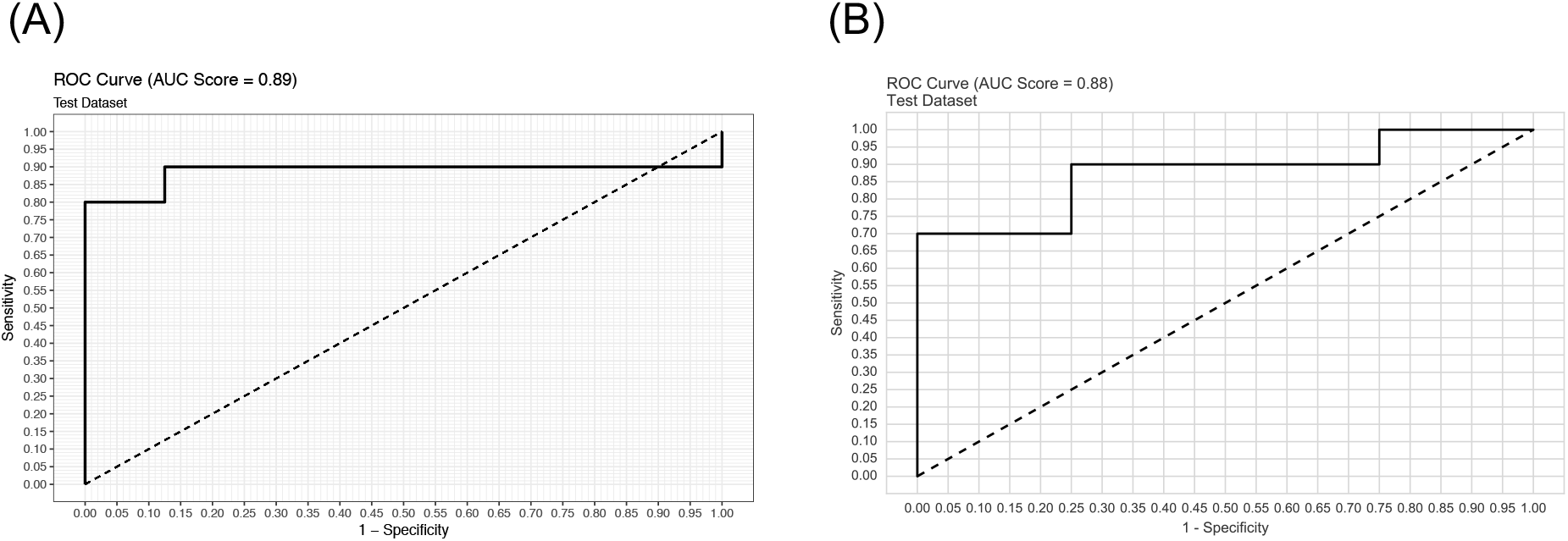
ROC Curve for the test dataset in (A) R and (B) Python.

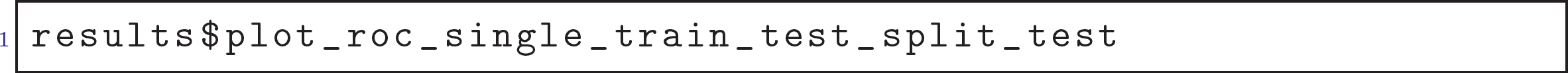

And in Python (see Figure 2B):

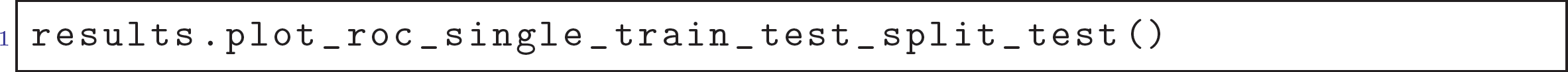

We can also examine the plot of the estimated beta coefficients. See Figure 3 where the coefficient means are represented by the dots and the error bars represent the standard deviations. We could reproduce the almost exact plot that appears in Ahn et al. (2016). These are true `ggplot2` objects in R and can be modified however needed:

**Figure 3:**
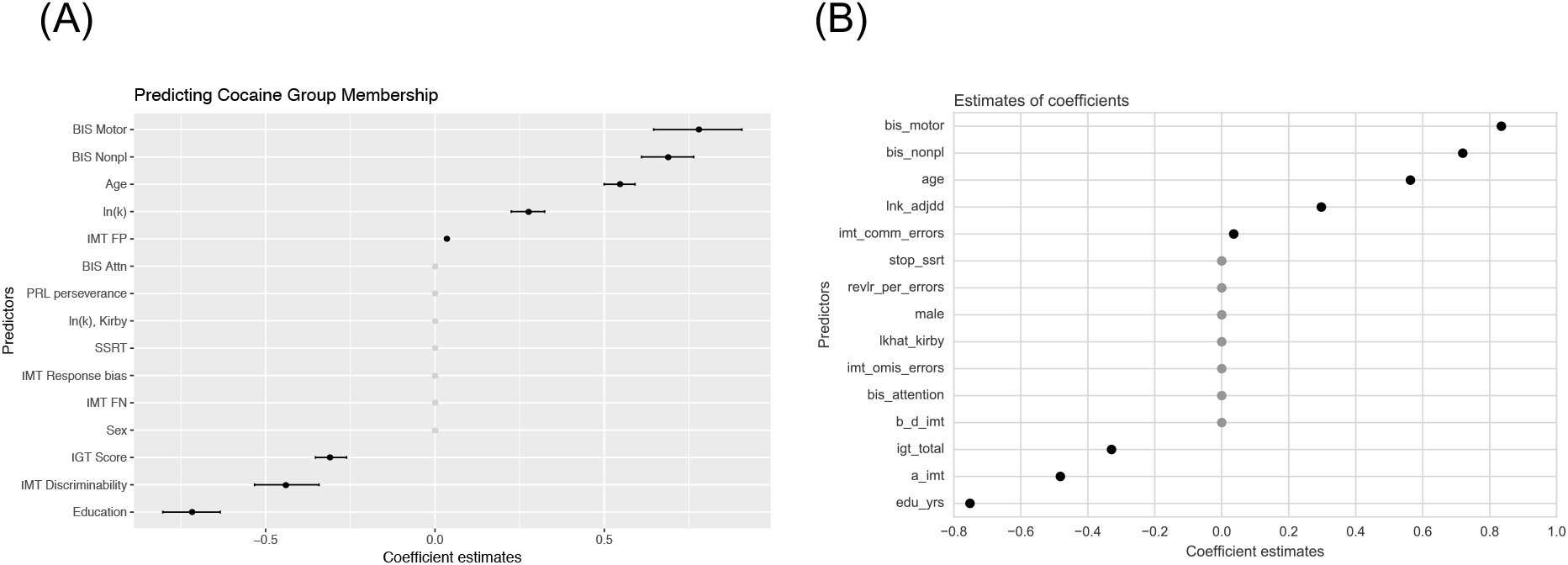
Multivariate patterns of impulsivity measures predicting cocaine dependence in (A) R and (B) Python. Error bars indicate 95% confidence intervals. See Ahn et al. (2016) for more details.

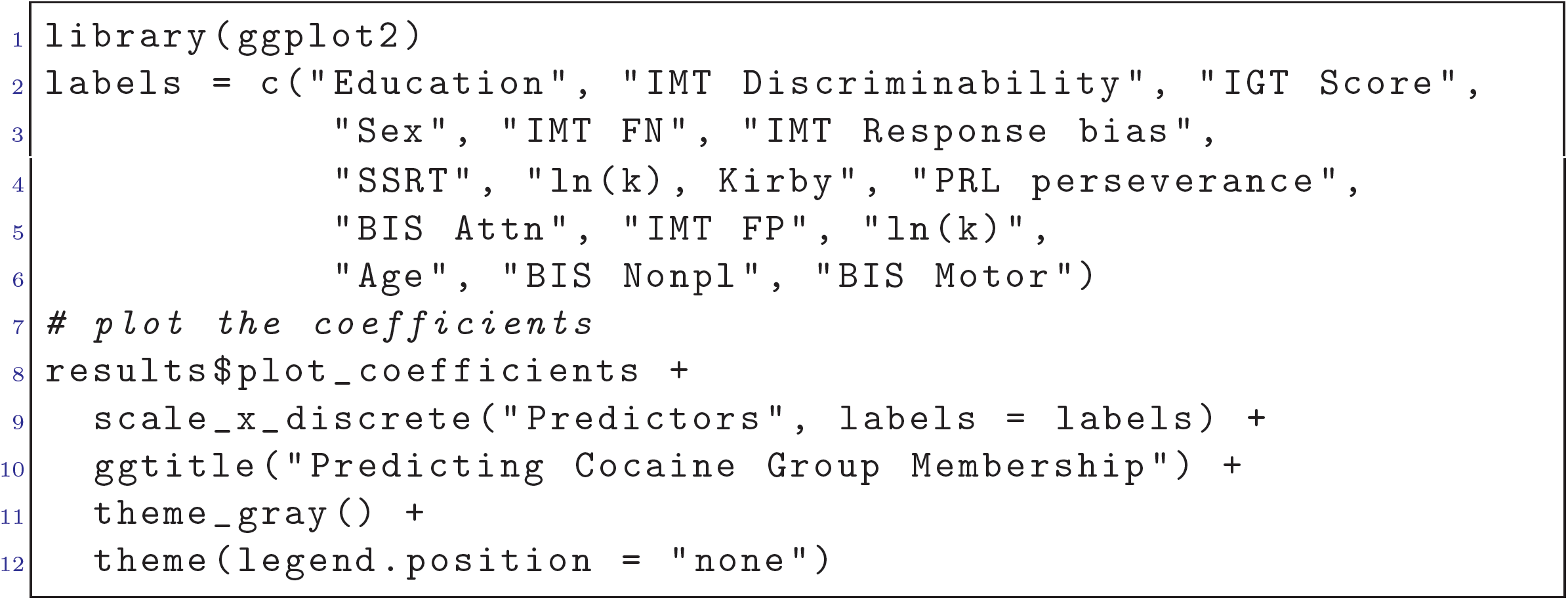

And in Python (see Figure 3):

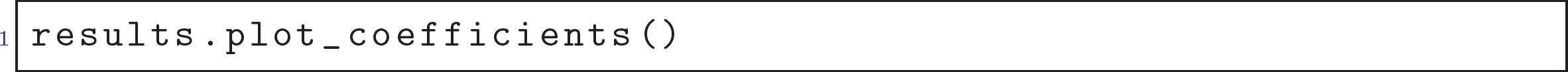

As seen in Figure 4A, by calling `results$plot_model_performance_train` in R, we can also examine the in-sample model performance generated for the train data set, which is the distribution of the AUCs of the ROC curves over 1,000 repetitions:

**Figure 4:**
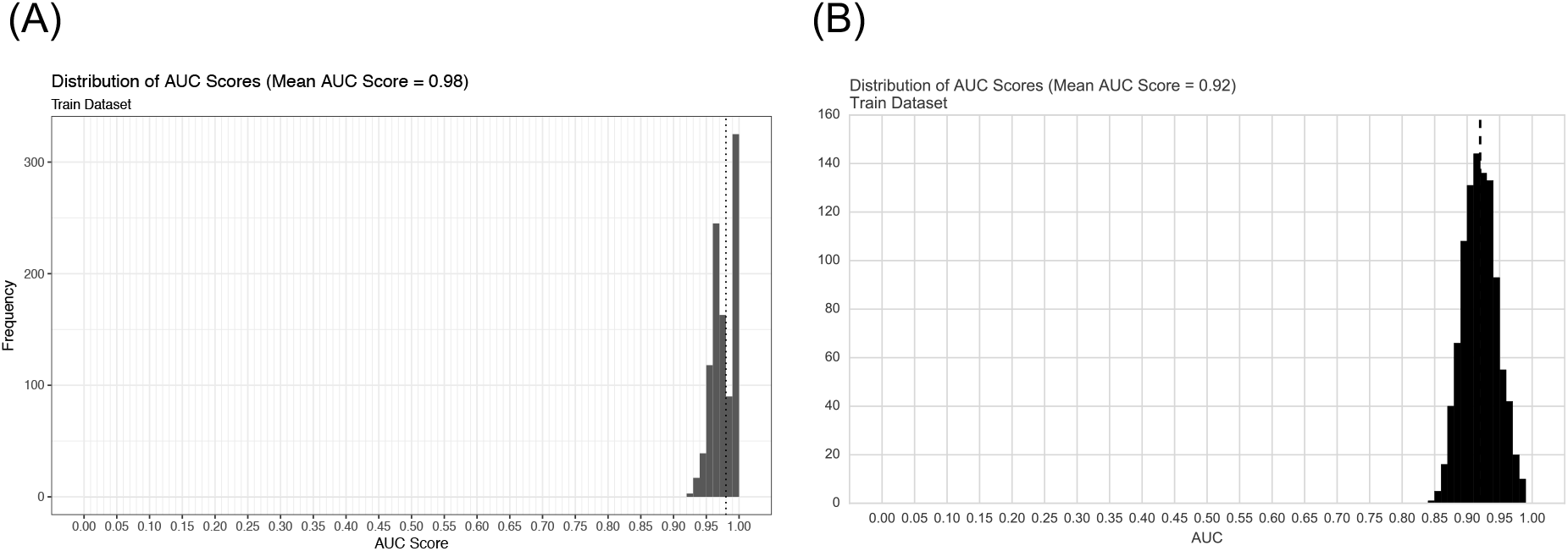
Measures of model performance for the train dataset in (A) R and (B) Python.

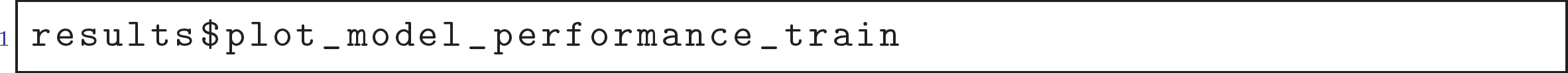

And in Python (see Figure 4B):

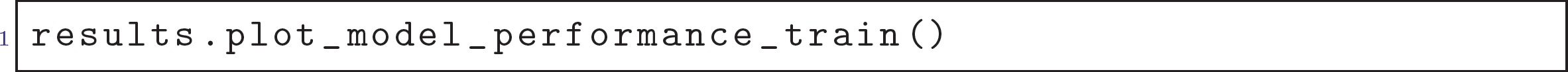

as well as the out-of-sample model performance generated for the test data set by calling `results$plot_model_performance_test` in R (see Figure 5A).

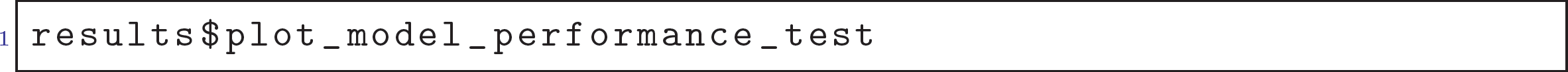

And in Python (see Figure 5B):

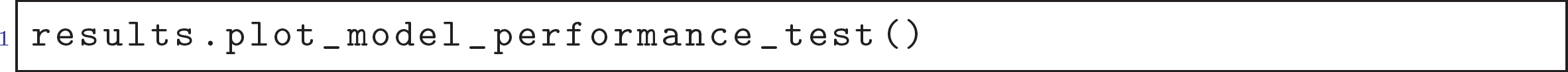

**Figure 5:**
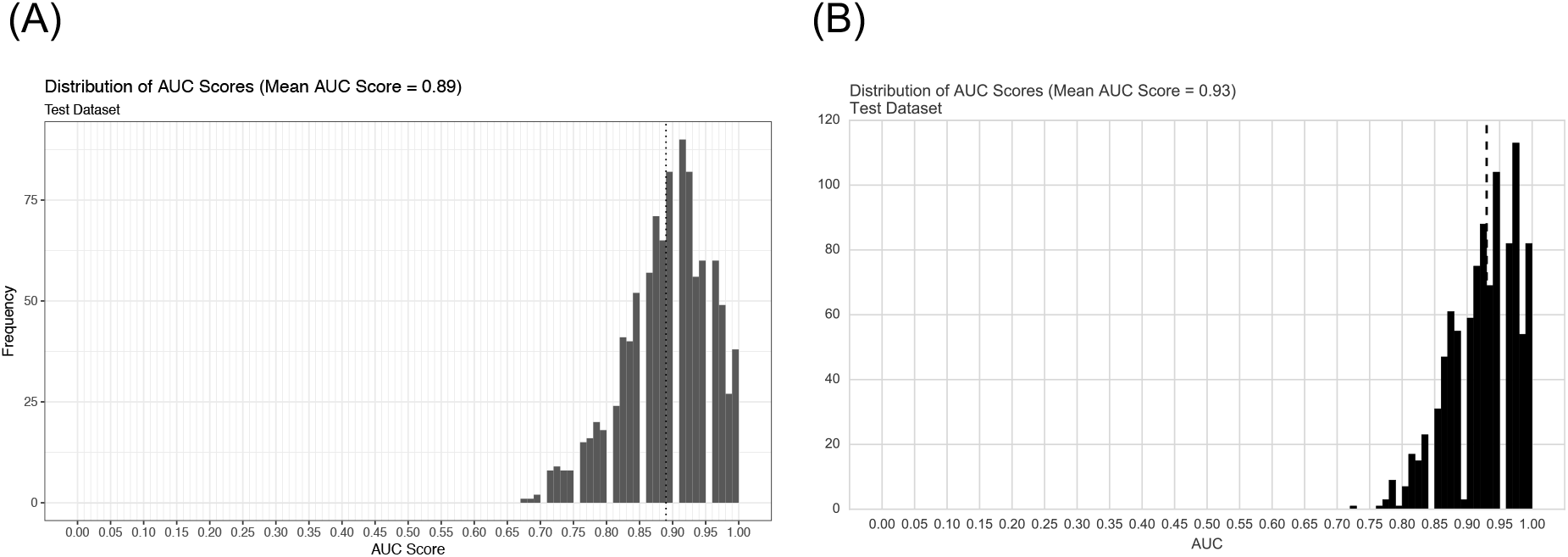
Measures of model performance for the test dataset in (A) R and (B) Python.

Users can run other algorithms as easily by following the same structured interface, with very few modifications to the parameters. For example, to run a random forest model, one would run in R:

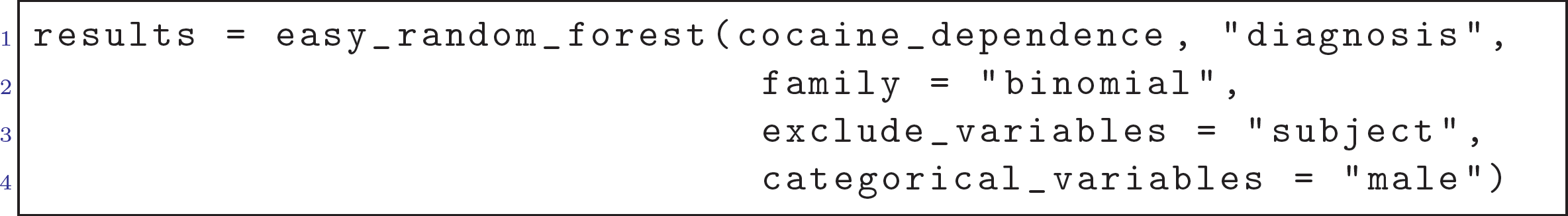

Or for a support vector machine model, one would run:

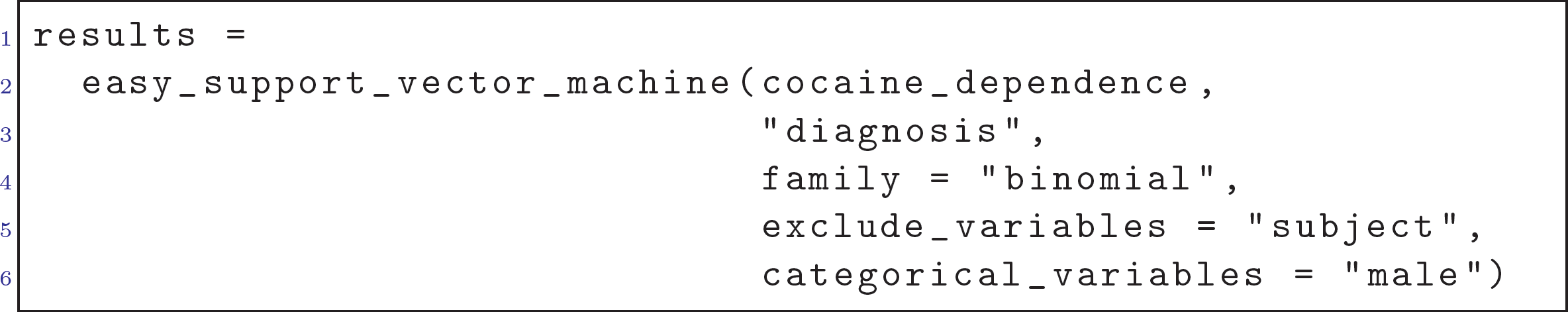

## 5. Implementing a New Algorithm

While penalized regression models, random forests, and support vector machines are among some of the most popular algorithms, an advanced user may wish to add an algorithm implemented elsewhere to `easyml` or perhaps even write their own algorithm. `easyml` makes this easy by allowing users to write `wrapper` functions to provide a common interface to those algorithms and pass wrapper functions into `easyml` functions. In Appendix A, we provide an example where we wrap an algorithm that uses averaging over several neural networks. The reader is referred to the `caret` (Kuhn, 2016) documentation for details on the `caret::avNNet` function.

## 6. Comparison to Similar Toolkits and Frameworks

R and Python both have a wide ecosystem of machine learning toolkits. `caret` and `mlr` (Bischl et al., 2016) are perhaps the most similar packages to `easyml` in R while `scikit-learn` (Pedregosa et al., 2011) is perhaps the most similar package to `easyml` in Python. Indeed these packages contain algorithms for regression and classification tasks, tools for preprocessing and model interpretation, and all focus on lowering the barrier to entry for machine learning for non-experts. Unlike other packages, with `easyml`, users can use standardized recipes for common machine learning techniques and produce journal-quality visualizations, all in a single line of coding.

## 7. Conclusions and Outlook

In conclusion, the `easyml` package fits a specialized niche, and further lowers the barrier to entry to machine learning. Practitioners have immediate access to powerful machine learning algorithms in a single-line of coding in R or Python, without worrying about their implementation or best practices for each algorithm. Researchers with strong programming skills can leverage the `easyml` library to provide customized extensions quickly. We also warn users that the use of `easyml` without sound understanding of machine learning can be potentially dangerous: we recommend users understand the basic concepts of machine learning and each algorithm they use. Otherwise they may incorrectly use algorithms implemented in `easyml` or interpret the results in a misleading way.

Next steps for `easyml` are likely to include more algorithms and additional recipes and convenience functions to further lower the barrier to entry for machine learning. We also plan make it easy to use `easyml` on neuroimaging data. Specifically, we will allow users to apply a machine learning algorithm to functional magnetic resonance imaging (fMRI) data and produce journal-quality brain maps in a single line of coding.

## 8. Author contributions

W.-Y.A. conceived the project. W.-Y.A., P.H., and N.H. programmed codes and designed/built the package.

## Appendix A.

Below we demonstrate an ordinary way of fitting the `avNNET` model of the `caret` package on the cocaine dataset in R. We see that while we can build this model relatively easily, it takes some extra work to build such as removing the first and second columns from the `cocaine_dependence` dataset. Furthermore, to evaluate this model multiple times with customized train-test splits, preprocessing data, visualize outputs, users need to program many lines of code additionally.

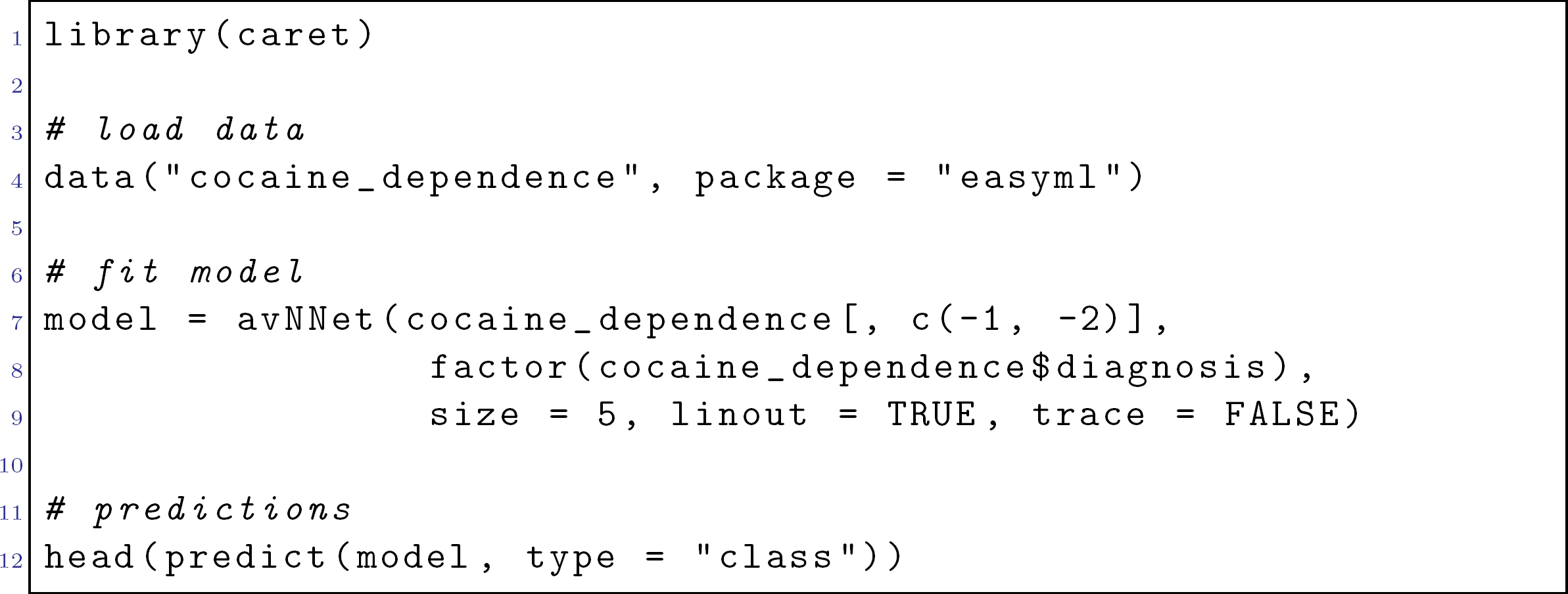

Here we demonstrate the `easyml` way of using this model (see below how we wrapped the algorithm into `easyml`). We see that with very few lines of code, we can enjoy all the features and benefits of `easyml`. For example, users can examine model performance by calling `b$plot_predictions_train_mean` and `$plot_metrics_test_mean`, etc.

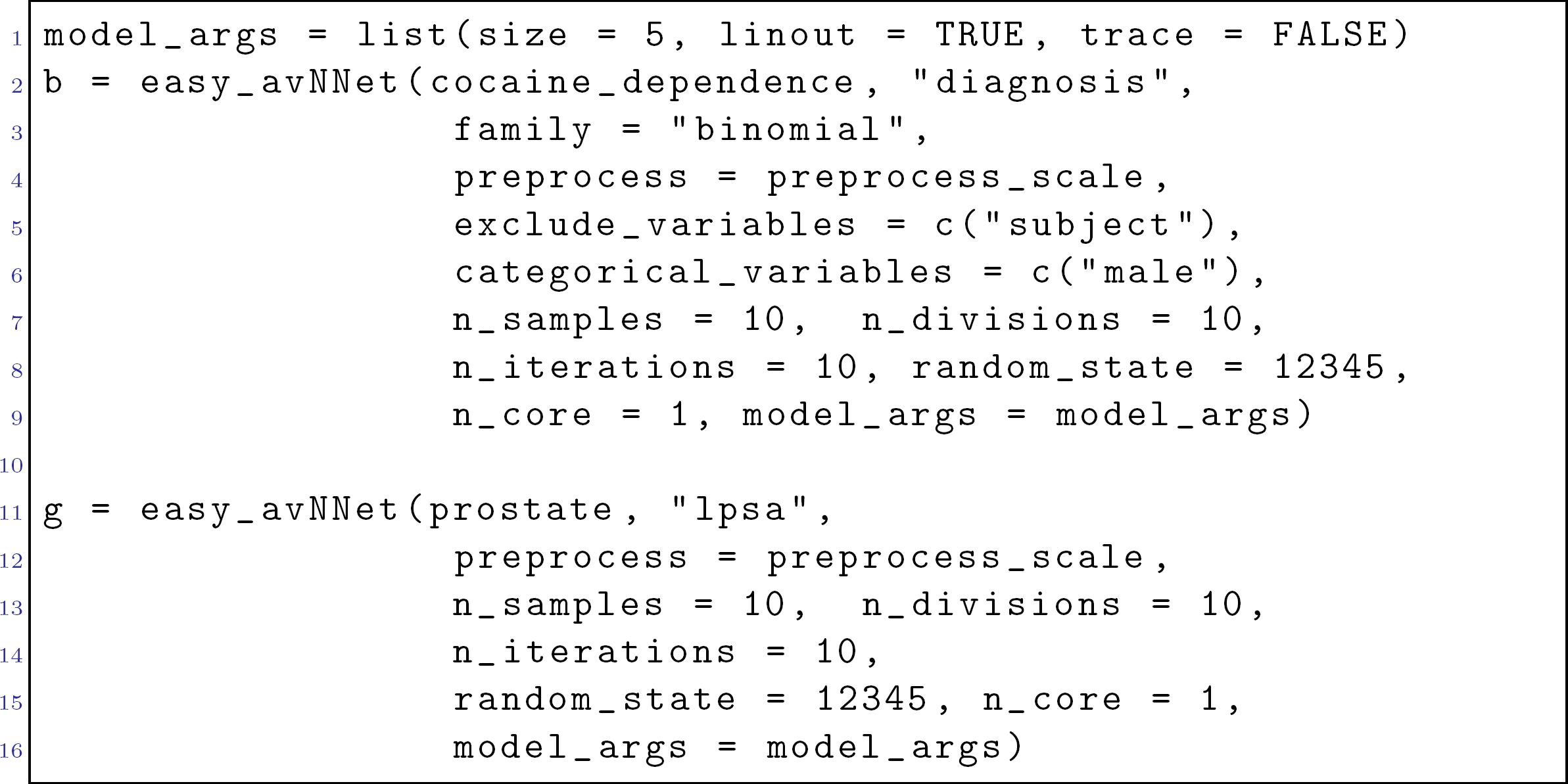

Here we show how we wrapped the `avNNet` algorithm into `easyml`.

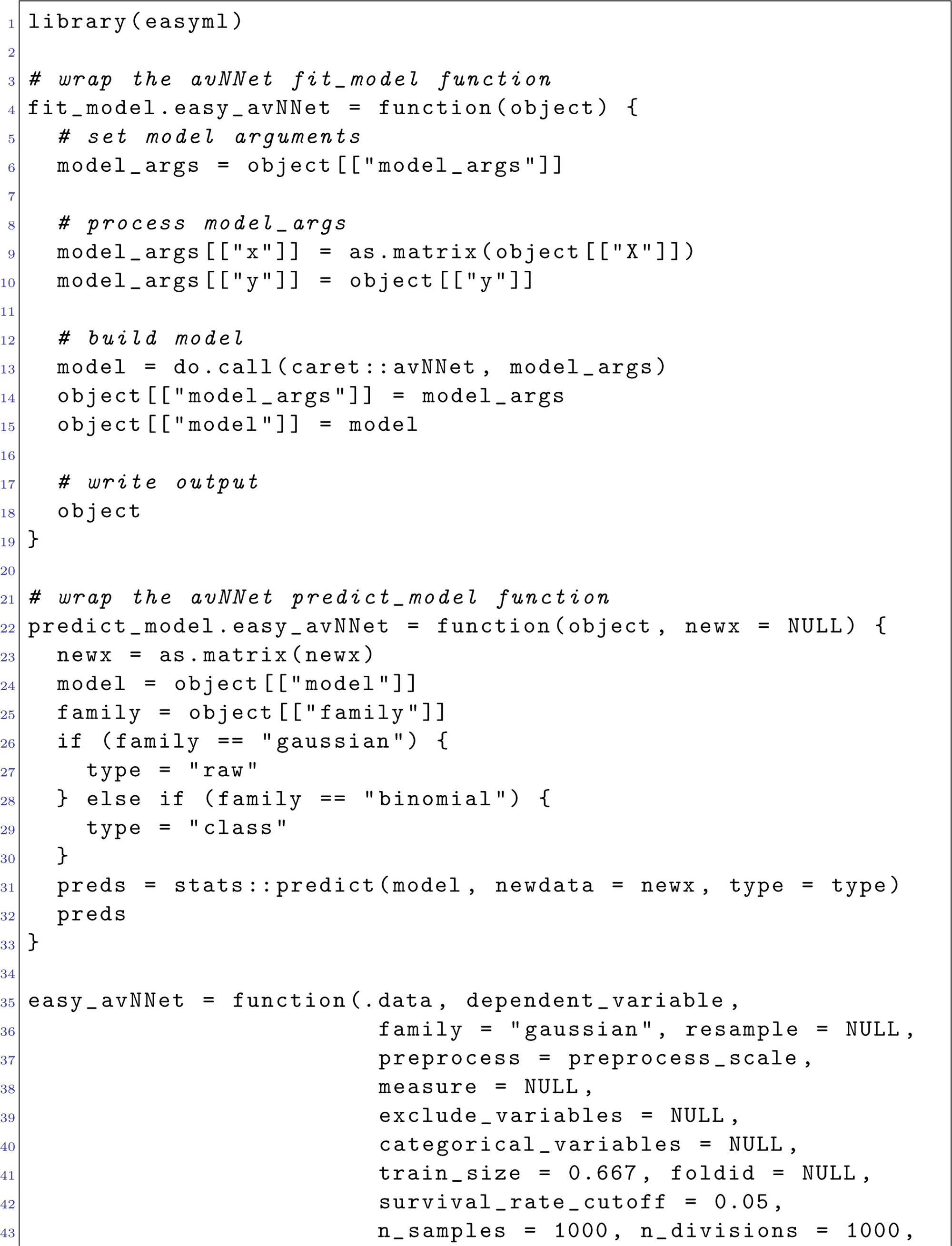

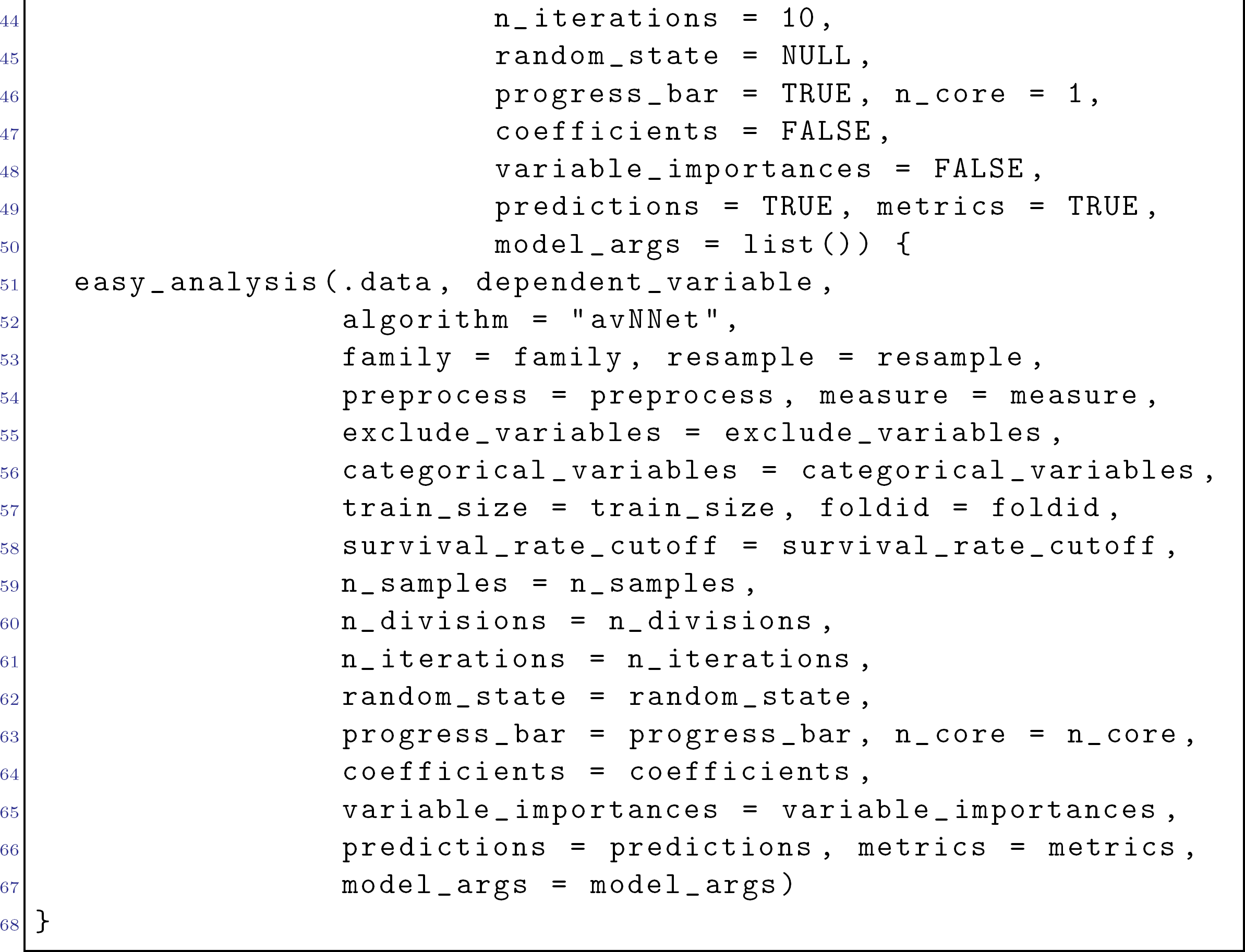

## References

Woo-Young Ahn and Jerome R Busemeyer. Challenges and promises for translating computational tools into clinical practice. Current Opinion in Behavioral Sciences, 11:1–7, 2016.

Woo-Young Ahn and Jasmin Vassileva. Machine-learning identifies substance-specific behavioral markers for opiate and stimulant dependence. Drug and alcohol dependence, 161: 247–257, 2016.

Woo-Young Ahn, Kenneth T Kishida, Xiaosi Gu, Terry Lohrenz, Ann Harvey, John R Alford, Kevin B Smith, Gideon Yaffe, John R Hibbing, Peter Dayan, et al. Nonpolitical images evoke neural predictors of political ideology. Current Biology, 24(22):2693–2699, 2014.

Woo-Young Ahn, Divya Ramesh, Frederick Gerard Moeller, and Jasmin Vassileva. Utility of machine-learning approaches to identify behavioral markers for substance use disorders: Impulsivity dimensions as predictors of current cocaine dependence. Frontiers in Psychiatry, 7, mar 2016. doi: 10.3389/fpsyt.2016.00034. URL https://doi.org/10.3389%2Ffpsyt.2016.00034.

Woo-Young Ahn, Nathaniel Haines, and Lei Zhang. Revealing neuro-computational mechanisms of reinforcement learning and decision-making with the hbayesdm package. Computational Psychiatry, 1(1), 2017.

Bernd Bischl, Michel Lang, Lars Kotthoff, Julia Schiffner, Jakob Richter, Erich Studerus, Giuseppe Casalicchio, and Zachary M. Jones. mlr: Machine learning in r. Journal of Machine Learning Research, 17(170):1–5, 2016. URL http://jmlr.org/papers/v17/15-066.html.

Leo Breiman. Random forests. Mach. Learn., 45(1):5–32, October 2001. ISSN 0885-6125. doi: 10.1023/A:1010933404324. URL http://dx.doi.org/10.1023/A:1010933404324.

Corinna Cortes and Vladimir Vapnik. Support-vector networks. Mach. Learn., 20(3):273–297, September 1995. ISSN 0885-6125. doi: 10.1023/A:1022627411411. URL http://dx.doi.org/10.1023/A:1022627411411.

Martin Drees. Implementierung und analyse von tiefen architekturen in r. Master’s thesis, Fachhochschule Dortmund, 2013.

Jerome Friedman, Trevor Hastie, and Robert Tibshirani. Regularization paths for generalized linear models via coordinate descent. Journal of Statistical Software, 33(1):1–22, 2010. URL http://www.jstatsoft.org/v33/i01/.

Nathaniel Haines, Matthew W. Southward, Paul Hendricks, Jeffrey F. Cohn, Jennifer S. Cheavens, and Woo-Young Ahn. Reading positive and negative emotion intensities from facial expressions using machine learning. in preparation.

Max Kuhn. caret: Classification and Regression Training, 2016. URL https://CRAN.R-project.org/package=caret. R package version 6.0-73.

Andy Liaw and Matthew Wiener. Classification and regression by randomforest. R news, 2(3):18–22, 2002.

Michael Lim and Trevor Hastie. Learning interactions via hierarchical group-lasso regularization. Journal of Computational and Graphical Statistics, 24(3):627–654, 2015.

David Meyer, Evgenia Dimitriadou, Kurt Hornik, Andreas Weingessel, and Friedrich Leisch. e1071: Misc Functions of the Department of Statistics, Probability Theory Group (Formerly: E1071), TU Wien, 2017. URL https://CRAN.R-project.org/package=e1071. R package version 1.6-8.

F. Pedregosa, G. Varoquaux, A. Gramfort, V. Michel, B. Thirion, O. Grisel, M. Blondel, P. Prettenhofer, R. Weiss, V. Dubourg, J. Vanderplas, A. Passos, D. Cournapeau, M. Brucher, M. Perrot, and E. Duchesnay. Scikit-learn: Machine learning in Python. Journal of Machine Learning Research, 12:2825–2830, 2011.

R Core Team. R: A Language and Environment for Statistical Computing. R Foundation for Statistical Computing, Vienna, Austria, 2016. URL https://www.R-project.org/.

Guido Rossum. Python reference manual. Technical report, Amsterdam, The Netherlands, The Netherlands, 1995.

Noah Simon, Jerome Friedman, Trevor Hastie, and Rob Tibshirani. Regularization paths for cox’s proportional hazards model via coordinate descent. Journal of Statistical Software, 39(5):1–13, 2011. URL http://www.jstatsoft.org/v39/i05/.

W. N. Venables and B. D. Ripley. Modern Applied Statistics with S. Springer, New York, fourth edition, 2002. URL http://www.stats.ox.ac.uk/pub/MASS4. ISBN 0-387-95457-0.

Iris Vilares, Michael J Wesley, Woo-Young Ahn, Richard J Bonnie, Morris Hoffman, Owen D Jones, Stephen J Morse, Gideon Yaffe, Terry Lohrenz, and P Read Montague. Predicting the knowledge-recklessness distinction in the human brain. Proceedings of the National Academy of Sciences, 114(12):3222–3227, 2017.

